# Corticotropin-releasing hormone in melanoma and non-melanoma skin cancer

**DOI:** 10.1101/2025.01.06.631434

**Authors:** Marcel Mueller, Susanne Melchers, Iris Mueller, Jochen Utikal, Julia Krug, Astrid Schmieder

## Abstract

**Background:** Corticotropin-releasing hormone (CRH) is involved in the regulation of immunological and cellular processes. Recently, CRH has been found to be expressed in skin cancers, where its expression appears to correlate with the degree of malignancy.

**Objective:** This study correlates CRH expression in melanoma metastases with patient survival and compares the intensity of CRH expression in melanoma to that in less aggressive skin cancer entities.

**Methods:** Tissue microarrays with cores from 94 melanomas and 40 melanocytic nevi and 51 slides from 41 basal cell carcinomas (BCC) and 10 squamous cell carcinomas (SCC) were immunohistochemically stained for CRH. The intensity of CRH expression in melanoma metastases was stratified by sex and correlated with patient survival. Furthermore, proliferation and apoptosis were assessed in CRH-stimulated A431 cells using enzyme-linked immunosorbent assays and an apoptosis detection kit.

**Results:** The intensity of CRH expression was higher in primary melanomas than in melanocytic nevi. Higher CRH expression was also found in melanoma metastases from women compared to men. However, higher CRH expression was correlated with reduced overall survival only in men. Compared to melanoma, BCCs and SCCs showed weaker CRH expression, which was in line with the finding that in vitro, CRH stimulation of the A431 cells reduced their proliferative activity.

**Conclusion:** CRH does not necessarily correlate with the degree of malignancy, as semi-malignant cancers such as BCC show higher levels of CRH expression than SCCs. In melanoma, CRH expression in metastases may be an important prognostic factor for overall survival in men, which needs further evaluation.

**Statement of contribution:** Marcel Mueller and Susanne Melchers performed all experiments and wrote the first draft of the manuscript. Iris Mueller helped with the experiments and reviewed the manuscript, Jochen Utikal assembled the tissue microarrays and reviewed the manuscript, Julia Krug helped with the figures and reviewed the manuscript, and Astrid Schmieder designed and supervised the project and corrected the manuscript.

## Introduction

The hypothalamic-pituitary-adrenal (HPA) axis is a complex endocrinological axis that maintains the body’s homeostasis. Corticotropin-releasing hormone (CRH) is a 41 amino acid long peptide that plays a central role in the HPA axis by stimulating Adrenocorticotropic hormone (ACTH) and ß-endorphin secretion and regulating glucocorticoid hormones. Outside the HPA axis, the highest concentrations of CRH are found in the heart, placenta, uterus, gastrointestinal tract, and adrenal gland (1). In 1995, Slominski et al. first identified the expression of CRH and its receptor in human skin, where CRH regulates peripheral homeostasis, including epidermal barrier function, pigmentation, and adnexal, dermal, and immune functions (2). CRH binds to CRH receptors 1 and 2 and induces local cortisol synthesis in diverse cells of the dermis (3). The hormone also promotes angiogenesis by stimulating mast cells, resulting in increased secretion of VEGF (4). Regarding proliferation, CRH favors keratinocyte differentiation and thus reduces keratinocyte proliferation (5). Since CRH has been described in several types of skin cancer, it is of interest to determine whether high CRH expression correlates with decreased or increased malignancy in human non-melanoma and melanoma skin cancers. Kim et al. found that only 10% of basal cell carcinomas (BCCs) showed strong immunoreactivity for CRH, whereas squamous cell carcinoma (SCC) showed strong expression in 70% and malignant melanoma in 80% of cases (6). Based on these data, the level of CRH expression appears to increase with the degree of malignancy, suggesting a tumorpromoting ability that may be partially mediated by its immunosuppressive function (6). The receptor for CRH, CRHR1, is expressed by keratinocytes and melanocytes, as well as their malignant counterparts, squamous carcinoma and melanoma cells. The effect of CRH on these cells is not fully understood and warrants further investigation. It is known that CRH induces a dose-dependent increase in intracellular calcium levels in melanoma cell lines and in the epidermoid carcinoma cell line A431 (1). In immortalized melanocytes, CRH has the ability to influence cell survival by inhibiting early and late apoptosis (7).

The aim of this study was to analyze CRH expression in nevi and compare it to primary melanoma and melanoma metastasis. The expression of CRH in melanoma metastases was further correlated with patient survival. In addition, CRH was evaluated in non-melanoma skin cancer and its effect on proliferation and apoptosis of A431 cells was analyzed in vitro.

## Methods

### Cell lines

The human epidermoid carcinoma cell line A431 (Merck, Darmstadt, Germany) was cultured in EMEM (30-2003™, ATCC®, Wesel, Germany) supplemented with 2 mM L-glutamin (Sigma-Aldrich, Munich, Germany), 1 % NEAA (Sigma-Aldrich, Munich, Germany), 10 % fetal calf serum (FCS, Biochrom, Berlin, Germany), 100 I.U./ml penicillin, and 100 µg/mL streptomycin (Pen/Strep, Biochrom, Berlin, Germany). The cell line was cultured at 37°C in a 5% CO_2_ enriched atmosphere.

### Human samples

51 formalin-fixed paraffin-embedded non-melanoma skin cancer specimens that were surgically removed or biopsied between 2011 and 2020 at the Department of Dermatology, Venerology, and Allergology, University of Heidelberg, Mannheim, Germany, were analyzed for this study.

Also, three Tissue-Microarrays (TMAs) with 49 malignant melanomas à two cores, 45 metastatic lesions from melanoma à two cores, and 40 melanocytic nevi from melanoma patients à two cores were stained. The study was performed according to federal laws and regulations as well as institutional policies of the Medical Faculty Mannheim. The local Ethics Committee granted ethical approval for all experiments performed with human patient samples (reference number: 2010-318N-MA and 2014-835R-MA). Informed consent was collected from all patients and the data was analyzed anonymously.

### Immunohistochemistry and light microscopy

Formalin-fixed, paraffin-embedded (FFPE) human tissue samples were deparaffinized using a descending xylene/alcohol series followed by heat-induced antigen retrieval (pH 6/9). Subsequently, the samples were treated with peroxidase blocking solution (Dako by Agilent, Waldbronn, Germany), blocked with 3 % BSA and incubated with anti-CRH antibody (ab8901, abcam, Cambridge, UK) in a humid chamber at 4°C overnight. Rabbit IgG (Cell Signaling Technology, Danvers, Massachusetts, USA) was used as an isotype control for the anti-CRH antibodies. After incubation with the appropriate HRP-conjugated secondary antibody (Dako by Agilent, Waldbronn, Germany) for 1 h at room temperature, AEC chromogen solution (Dako by Agilent, Waldbronn, Germany) was applied for visualization. 10% Mayer’s Haemalaun (Merck, Darmstadt, Germany) was used for counterstaining. Images were captured with a Leica DCRE microscope, Leica DC500 camera, and software system (Leica, Wetzlar, Germany).

The degree of CRH-expression in non-melanoma skin cancer was graded as follows: 0, negative; +, weakly positive; ++, strongly positive; +++, very strongly positive. Three dermatologists independently evaluated the results.

The Tissue-Microarrays (TMAs) were stained according to the same protocol. The cores were then scored from 0 to 100 according to the number of positive cells (Figure 1).

**Figure 1.**
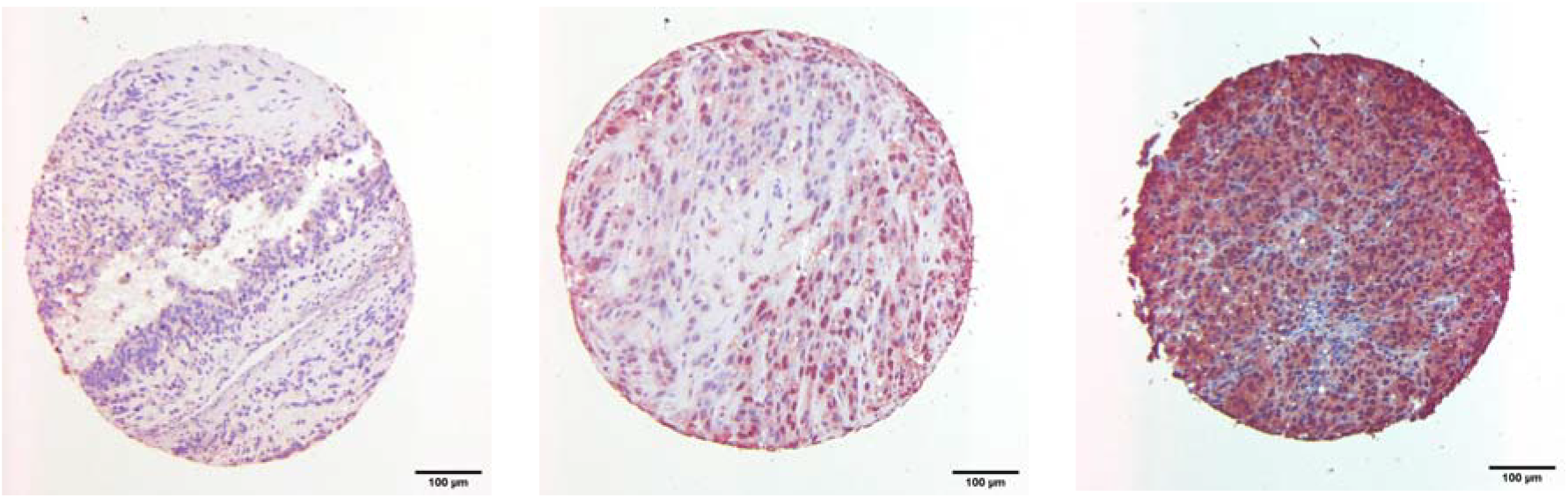
CRH expression in human melanocytic lesions. Three exemplary stains of three TMA cores composed of human primary melanoma specimens (A), human melanoma metastatic lesions (B), and melanocytic nevus derived from melanoma patients (C) for CRH with a score of 20 (A), 50 (B), and 90 (C). The samples were evaluated and a score from 0 to 100 was assigned.

### Enzyme-linked immunosorbent assay (ELISA)

1 × 106 A431 cells were seeded in 2 mL EMEM complete and treated with 0 nM, 1 nM, 10 nM, or 100 nM CRH (Sigma-Aldrich, Munich, Germany) for 24 h. Cell supernatants were collected and centrifuged at 1000 x g for 10 min to obtain cell-free supernatants. ELISA was performed according to the manufacturer’s instructions (human IL-8/CXCL8 DuoSet ELISA, R&D Systems, Wiesbaden, Germany). The relative absorbance was measured at a wavelength of 450 nm, and the IL-8/CXCL8 concentrations were calculated using a standard curve. Each supernatant was analyzed in triplicate.

### Proliferation assay

For the proliferation assay, 2 × 105 A431 cells were seeded per well in 96-well plates. After the cells became adherent, the medium was replaced with EMEM complete containing 0, 1, 10, or 100 nM corticotropin-releasing hormone (CRH) for 24 h. To quantify the cell proliferation, BrdU was added, which was incorporated into the DNA of dividing cells. The assay was performed according to the manufacturer’s instructions (ab126556, BrdU Cell Proliferation Kit ELISA (colorimetric), Abcam, Cambridge, UK). Absorbance was read at 450 nm with a reference wavelength of 550 nm using a microplate reader. Each concentration was analyzed in a sextet.

### Apoptosis assay

For apoptosis detection, 2 × 105 A431 cells were seeded per well in 6-well plates. After the cells became adherent, the medium was replaced with EMEM containing 0, 1, 10 or 100 nM CRH. After incubation for 24 hours, the cells were gently trypsinized and washed once with serum containing medium. Prior to annexin staining, cells were centrifuged and excess media was decanted at the beginning of processing. 1 × 105 cells were stained with Annexin V-FITC conjugated antibody and propidium iodide from the Annexin V-FITC apoptosis detection kit (ab14085, Abcam, Cambridge, UK) according to the manufacturer’s instructions. Annexin V-FITC binding was read on the flow cytometer.

### Statistics

Statistical analyses of all data were calculated by using GraphPad Prism 7.0 (GraphPad Software, USA). Statistical significance was assessed by using Student’s t-test or by one-way ANOVA and Bonferroni. Not normally distributed data were analyzed with Kruskal-Wallis-Test and Mann-Whitney-Test. The level of significance is indicated by asterisks (^***^ ≤ 0.001; ^**^ ≤ 0.01 and ^*^ ≤ 0.05). Error bars depict the standard error of the mean (SEM) of each experiment.

## Results

### High CRH expression in melanoma metastases in men, but not in women, correlates negatively with survival

Three TMAs with samples from 49 primary melanomas, 45 metastatic melanomas and 40 melanocytic nevi from melanoma patients were stained for CRH expression. The staining was scored from 0 to 100 according to the number and intensity of positive cells (Figure 1). Quantification revealed a significantly higher expression of CRH in primary melanomas than in nevi. Melanoma metastases also showed higher CRH expression than nevi, although the difference was not significant (Figure 2A). For CRH expression, a significant difference between female and male metastatic melanomas was evaluated. Females had a significantly higher level of CRH expression than males (Figure 2B). Survival data of male and female patients with metastatic melanoma were correlated with CRH expression. In males, there was a correlation between survival and CRH expression, with high CRH expression associated with lower survival (Figure 2C).

**Figure 2.**
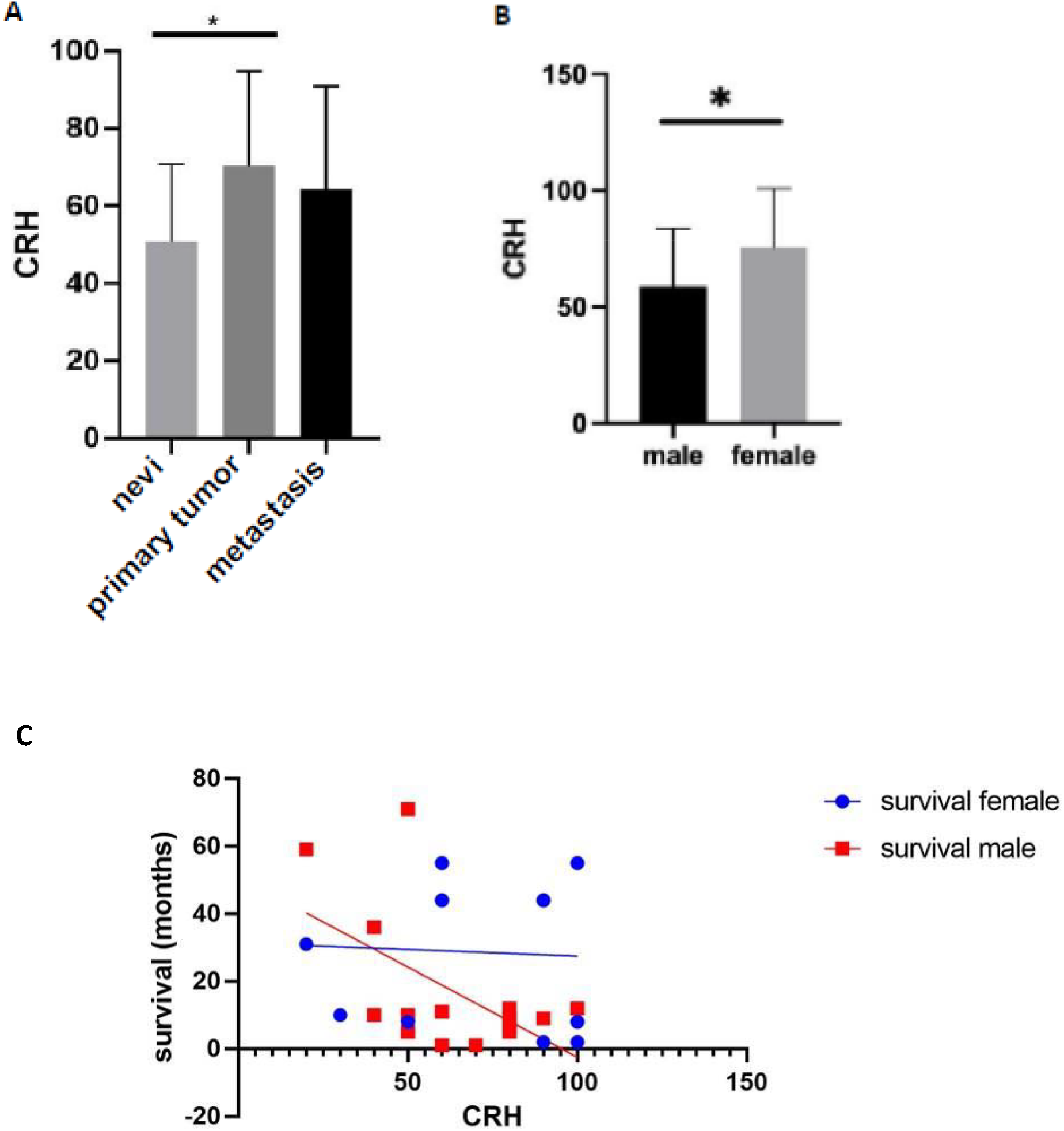
CRH expression is increased in primary melanomas compared to melanocytic nevi and it correlates with a decreased survival in men, but not in women. CRH expression in melanocytic nevi, primary melanomas and melanoma metastases assessed by quantification of immunohistochemical staining of three TMAs (A). Sex differences in CRH expression in metastatic melanoma specimens assessed by quantification of immunohistochemical staining of three TMAs (B) and correlation of these data with the overall survival rate in men and women (C).

### Non-melanoma skin cancers show a weaker CRH expression than melanoma

Fifty-one samples of various basal cell carcinomas (BCCs) and squamous cell carcinomas (SCCs) were immunohistochemically stained for CRH expression to determine the expression of CRH in human non-melanoma skin cancers (Figure 3). The expression pattern was scored as negative (0), weakly positive (+), strongly positive (++), very strongly positive (+++). Compared to the staining intensity observed in melanoma, the staining was generally weaker. Only some nodular (7 of 15) and superficial BCCs (7 of 13) showed strong or very strong CRH expression (Figure 3E).

**Figure 3.**
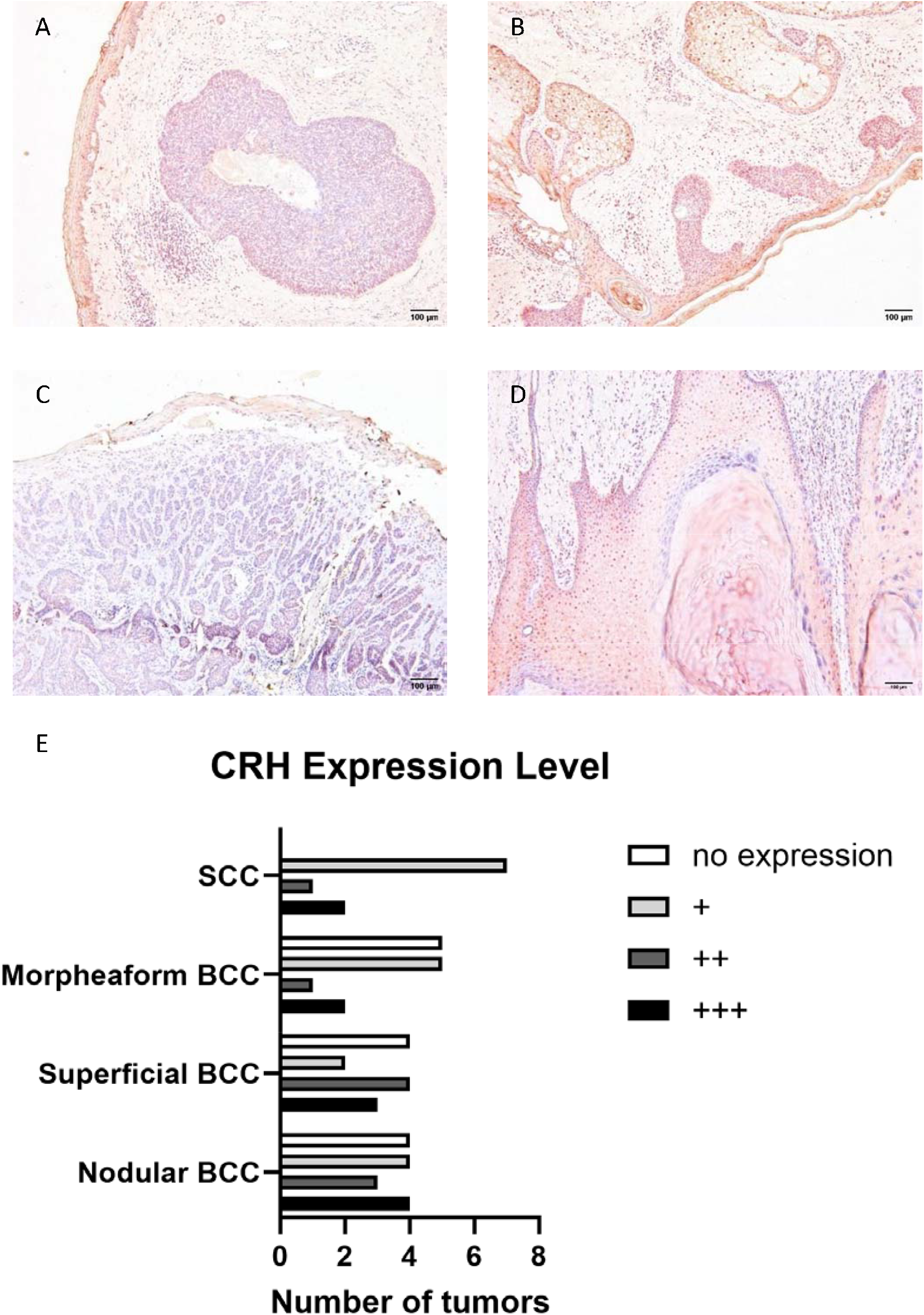
CRH expression in human basal cell carcinomas (BCC), and squamous cell carcinomas (SCC). Representative images of immunohistochemical stainings for CRH in nodular BCC (A), superficial BCC (B), morpheaform BCC (C), and SCC (D). Scale bar = 100 μm. (E) Number of tumors with no, weakly positive, strongly positive or very strongly positive CRH expression (quantification of immunohistochemical stainings). The expression pattern was graded as follows: 0 = negative, + = weakly positive, ++ = strongly positive, +++ = very strongly positive. Each staining was evaluated independently by three dermatologists and the mean was calculated.

### CRH inhibits proliferation and reduces late apoptosis in A431 cells

To determine whether CRH exerts a functional effect on non-melanoma skin cancer cells, as has been described for various human skin cells (Slominski et al 2005), we performed functional in vitro assays using the epidermoid carcinoma cell line A431. Since CRH has been described to modulate IL-8 secretion in various cells, we treated A431 cells with 1 nM, 10 nM, or 100 nM CRH for 24 hours and measured IL-8 levels in cell supernatants by ELISA. Notably, 1 nM CRH was able to significantly increase IL-8 secretion from A431 cells, whereas higher concentrations of CRH abolished this effect (Figure 4A).

**Figure 4.**
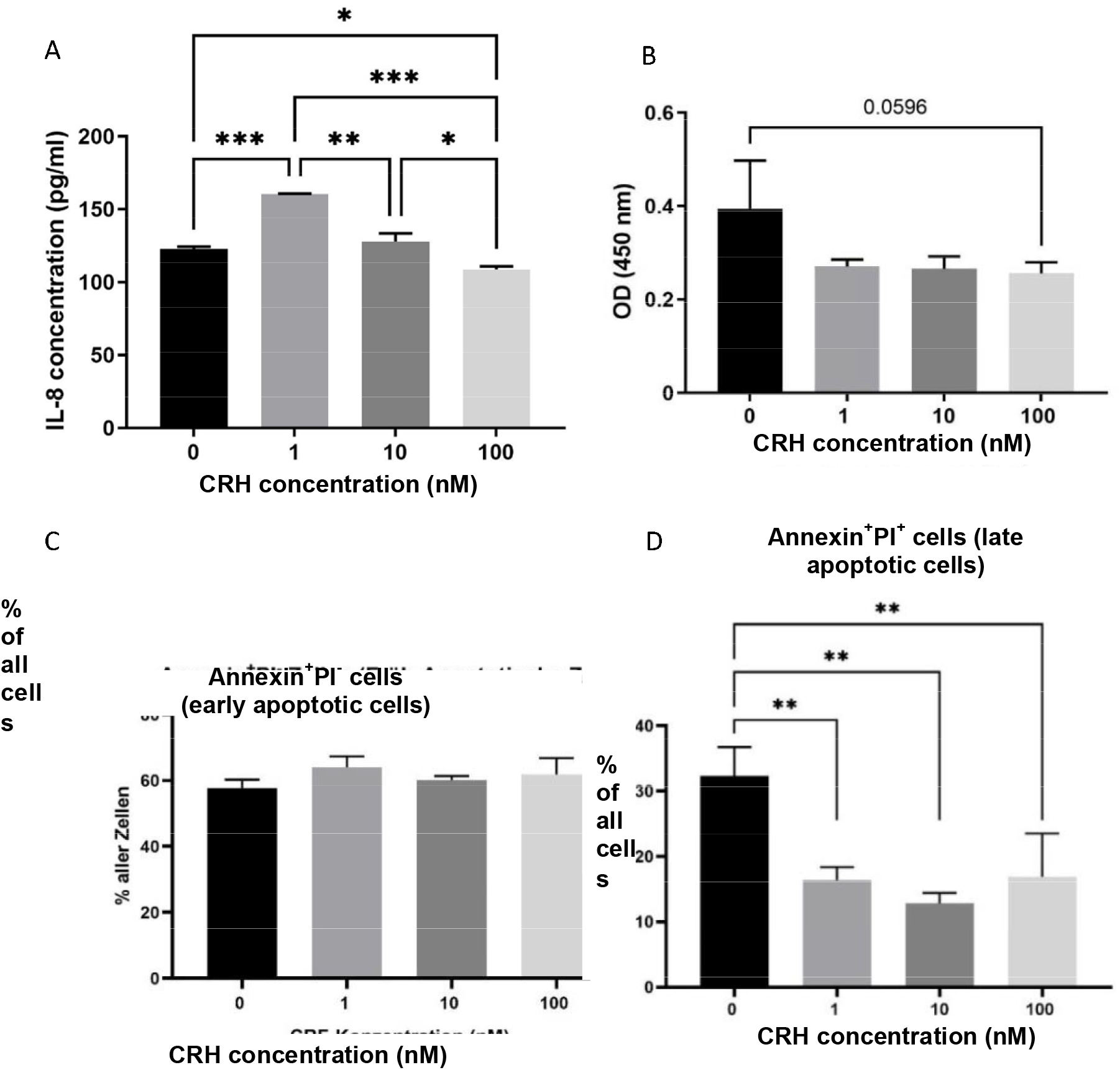
In A431 cells, CRH at 1 nM increases IL-8 secretion and inhibits both proliferation and apoptosis. IL-8 secretion of A431 cells after treatment with 1 nM, 10 nM, or 100 nM CRH or 24 hours as assessed by enzyme-linked immunosorbent assay (A). Proliferation of A431 cells after treatment with 1 nM, 10 nM, or 100 nM CRH as measured by using the BrdU cell proliferation assay (B). Apoptosis of A431 after treatment with 1 nM, 10 nM, or 100 nM CRH using the Annexin V-FITC apoptosis assay. Early apoptotic cells (C) and late apoptotic cells (D) are shown. Data are presented as mean ± SD. ^*^p < 0.05, **p < 0.01, ^***^p < 0.001.

Next, the effects on proliferation of A431 cells after treatment with CRH were analyzed. There was a trend towards a lower proliferation rate in A431 cells treated with CRH (Figure 4B). However, the difference did not reach significance.

Based on published data, CRH seems to have an effect on early and late apoptosis. Therefore, we additionally investigated the effect of CRH on cell death. In the apoptosis assay, there was no difference in early apoptotic cells (Figure 4C). However, a significant effect was observed in late apoptotic cells. The proportion of apoptotic A431 cells after CRH stimulation was significantly lower than that of untreated cells (Figure 4D).

## Discussion

In this study, we demonstrated that primary melanomas exhibit significantly higher expression of CRH compared to benign melanocytic nevi. Additionally, we observed that increased CRH expression is more pronounced in melanoma metastases from women than from men. Interestingly, higher CRH expression in melanoma metastases correlated with worse overall survival only in men. The higher CRH expression in melanoma is consistent with the findings of Kim et al. who also described a significant increase in CRH expression in primary melanoma compared to nevi (6). The higher CRH expression in melanoma metastases from females has not been described yet.

In 1993, Vamvakopoulos et al. identified five palindromic estrogen response elements (ERE) in the 5’ flanking region of the human CRH gene (8). In following publications, the group proposed a direct ER-mediated stimulation of CRH synthesis and secretion as a potential cause for the sexual dimorphism of the human stress response (9). Regarding androgens, a specific androgen-responsive element (ARE) was identified in the human CRH promotor region. Cotransfection experiments revealed that the AR in the presence of testosterone can repress CRH promotor activity through the ARE site (10). Thus, androgens can inhibit human CRH production directly through the androgen receptor. This could potentially explain the increased CRH expression observed in melanoma metastases of females.

Explaining the decreased survival of men with high CRH-expressing metastatic melanoma is not straightforward. However, it is known that estrogen-responsive melanomas are associated with the superficial spreading melanoma subtype, which generally has a better prognosis compared to the nodular form. Furthermore, female melanomas tend to metastasize more slowly than male melanomas, and their pattern of metastatic spread differs, often resulting in more locoregional recurrences in women (11).

CRH expression has been found in other cancer types such as breast, ovarian, and endometrial cancers. There, CRH and its receptor expression has been correlated with a higher FIGO (Fédération Internationale de Gynécologie et d’Obstétrique) stage, probably due to increased CRH-induced Fas ligand mediated local T-cell apoptosis, thus facilitating tumor immunoescape (12, 13). In 1999, Arbiser et al. transfected the human epithelial cell line 293 with human CRH and implanted the transfected cells subcutaneously in nude mice. The CRH-bearing tumors showed a significant increase in microvessel density. In addition, stimulation with CRH resulted in increased migration of capillary endothelial cells (14). Therefore, a potential explanation for the increased mortality rate among men with increased CRH expression in melanoma metastasis could be attributed to the absence of protective estrogen effects.

Additionally, the increased locoregional immunosuppressive and proangiogenic effects of CRH might contribute to the development of a more aggressive melanoma subtype. Kim et al. conducted an immunohistochemical analysis of CRH, ACTH, and α-MSH expression in benign, semi-malignant, and malignant skin tumors. They found strong CRH expression in melanoma and SCC, and intermediate in BCC. Consequently, they concluded that CRH expression increases with the degree of malignancy (6). To validate this finding we stained 51 different BCCs and SCCs and scored the expression intensity. Quantification showed the highest CRH expression in the semi-malignant superficial and nodular BCCs, and not in SCCs, thus failing to confirm the association with a more aggressive tumor phenotype.

It is known that CRH promotes the survival of melanocytes during starvation and prevents cell proliferation (7). Furthermore, CRH seems to induce keratinocyte differentiation (5). We therefore analyzed the effect of CRH on the proliferation and apoptosis of A431 carcinoma cells *in vitro* and confirmed that CRH reduces their proliferative activity and late apoptosis, two cellular characteristics typically found in differentiation processes. Therefore, we believe that at least in non-melanoma skin cancer, CRH does not promote tumor growth per se, but rather exerts its effects on the tumor stroma by increasing tumor vasculature and reducing an effective immune response. These two effects may promote melanoma progression, but perhaps not so much non-melanoma skin cancer progression, as their differentiation status may correlate more with their aggressiveness. However, this contention needs to be evaluated in a larger cohort of patients.

One limitation of the present study is its retrospective nature. In addition, while we analyzed nearly 94 melanomas/melanoma metastases, we only quantified CRH expression of small tumor sections rather than the entire tumor. Additionally, we did not account for other factors crucial for survival prognosis, such as tumor thickness at initial diagnosis, lactate dehydrogenase levels, or the analysis of immune infiltrate.

## Conclusion

Semi-malignant tumors such as basal cell carcinoma exhibit higher levels of CRH expression compared to squamous cell carcinoma, suggesting that CRH may not correlate with malignancy in non-melanoma skin cancer. This is different from the pattern observed in melanoma, where benign nevi exhibit lower CRH expression than primary melanomas. Considering the correlation between high CRT expression an overall survival in men, CRH can be evaluated as a potential therapeutic target in this patient group.

## Acknowledgements

We thank Sayran Arif-Said, and Hilltrud Schönhaber for excellent technical assistance. This work was funded by the Deutsche Forschungsgemeinschaft (DFG, German Research Foundation) – Project number 259332240/RTG 2099 to A.S. an J.U.

## References

1. Slominski A, Wortsman J, Pisarchik A, Zbytek B, Linton EA, Mazurkiewicz JE, et al. Cutaneous expression of corticotropin-releasing hormone (CRH), urocortin, and CRH receptors. FASEB J. 2001;15(10):1678–93.

2. Slominski A, Ermak G, Hwang J, Chakraborty A, Mazurkiewicz JE, Mihm M. Proopiomelanocortin, corticotropin releasing hormone and corticotropin releasing hormone receptor genes are expressed in human skin. FEBS Lett. 1995;374(1):113–6.

3. Slominski AT, Zmijewski MA, Zbytek B, Tobin DJ, Theoharides TC, Rivier J. Key role of CRF in the skin stress response system. Endocr Rev. 2013;34(6):827–84.

4. Cao J, Papadopoulou N, Kempuraj D, Boucher WS, Sugimoto K, Cetrulo CL, et al. Human mast cells express corticotropin-releasing hormone (CRH) receptors and CRH leads to selective secretion of vascular endothelial growth factor. J Immunol. 2005;174(12):7665–75.

5. Zbytek B, Slominski AT. Corticotropin-releasing hormone induces keratinocyte differentiation in the adult human epidermis. J Cell Physiol. 2005;203(1):118–26.

6. Kim MH, Cho D, Kim HJ, Chong SJ, Lee KH, Yu DS, et al. Investigation of the corticotropin-releasing hormone-proopiomelanocortin axis in various skin tumours. Br J Dermatol. 2006;155(5):910–5.

7. Slominski A, Zbytek B, Pisarchik A, Slominski RM, Zmijewski MA, Wortsman J. CRH functions as a growth factor/cytokine in the skin. J Cell Physiol. 2006;206(3):780–91.

8. Vamvakopoulos NC, Chrousos GP. Evidence of direct estrogenic regulation of human corticotropin-releasing hormone gene expression. Potential implications for the sexual dimophism of the stress response and immune/inflammatory reaction. J Clin Invest. 1993;92(4):1896–902.

9. Torpy DJ, Papanicolaou DA, Chrousos GP. Sexual dimorphism of the human stress response may be due to estradiol-mediated stimulation of hypothalamic corticotropin-releasing hormone synthesis. J Clin Endocrinol Metab. 1997;82(3):982.

10. Bao AM, Fischer DF, Wu YH, Hol EM, Balesar R, Unmehopa UA, et al. A direct androgenic involvement in the expression of human corticotropin-releasing hormone. Mol Psychiatry. 2006;11(6):567–76.

11. Mervic L. Time course and pattern of metastasis of cutaneous melanoma differ between men and women. PLoS One. 2012;7(3):e32955.

12. Minas V, Rolaki A, Kalantaridou SN, Sidiropoulos J, Mitrou S, Petsas G, et al. Intratumoral CRH modulates immuno-escape of ovarian cancer cells through FasL regulation. Br J Cancer. 2007;97(5):637–45.

13. Dimas A, Margioula-Siarkou C, Politi A, Sotiriadis A, Papanikolaou A, Dinas K, et al. The expression and possible role of corticotropin-releasing hormone family peptides and their corresponding receptors in gynaecological malignancies and premalignant conditions: a systematic review. Prz Menopauzalny. 2023;22(4):227–35.

14. Arbiser JL, Karalis K, Viswanathan A, Koike C, Anand-Apte B, Flynn E, et al. Corticotropin-releasing hormone stimulates angiogenesis and epithelial tumor growth in the skin. J Invest Dermatol. 1999;113(5):838–42.

